# Biochemical and structural characterisation of MprF homologue, LpiA from *Agrobacterium*

**DOI:** 10.64898/2026.04.24.720753

**Authors:** Shardul Dhole, Pallavi Sabharwal, Kutti R. Vinothkumar

## Abstract

Multiple peptide resistance factor (MprF) are bi-functional enzymes encoded by several bacterial species and carry out the transfer of an amino acid from a charged tRNA to the lipid head group and further translocate the lipid across the membrane. Biochemical studies have revealed that the soluble synthase domain generates specificity and the structures of MprF have defined the general architecture of these enzymes, and that they can exist in different oligomeric states. Here, we characterise the gene product of lpiA, a MprF homologue from *Agrobacterium fabrum (*formerly called *A. tumefaciens* strain C58), a microbe that is commonly used in plant molecular biology. Cryo-EM analysis of AfMprF reveals a dimeric structure both in detergent micelle and in lipid nanodisc, and similar in architecture to the homologous enzyme from related Rhizobium sp. We further analyse some conserved residues in the soluble domain and suggest that the sulphur-aromatic motifs play a key role in substrate binding. Similar architecture of enzymes in closely related bacterial species of Agrobacterium and Rhizobium hints an evolutionary relationship but the importance of these oligomeric states in vivo remains to be analysed.

## Introduction

We have been interested in understanding the mechanism of the Multiple peptide resistance factor (MprF), a bi-functional enzyme that largely exists in bacteria. MprF has two domains, a transferase domain that uses aminoacylated tRNA as a substrate and modifies the lipid head group with an amino acid and the transmembrane domain (TMD), which flips the lipid from one leaflet to the other^1–5^. The modified lipids change the property of the membrane, and allow the bacteria to cope with different environmental conditions and also confer resistance against cationic antimicrobial peptides (CAMPs)^1,3,6,7^. The soluble synthase (or transferase) domain of MprF is well characterised revealing two GCN5-related N-acetyltransferase (GNAT) folds and potential sites for substrate binding ^8,9^. Previous studies have shown that the soluble domain determines the substrate specificity and it can modify the lipids as a separate entity^9,10^. Replacement of the soluble domain between homologues results in differential modifications of the lipids^10^. Characterisation of MprF from different organisms has revealed some enzymes to be extremely specific while others are promiscuous ^2,11–13^. In some organisms, more than one MprF gene is found with differing specificity and in *Enterococcus faecalis*, only one of the MprF (MprF2) has shown to be active, while no activity has been observed so far for MprF1^14^.

The recent structure determination of the full-length structures of MprF by cryo-EM from different species has shed light on the relative positions of the soluble and the membrane domain and the oligomeric states^10,15,16^. The membrane domain has 14 transmembrane (TM) helices including two re-entrant helices similar to those found in transporters^17^, with TM14 connecting the soluble domain. The structure from Rhizobium species revealed a dimer with the interface formed by TM helices 4, 5, 6 and 7a^10^. In contrast, MprF from *Pseudomonas aeruginosa* has been determined as a monomer or as a dimer that is distinct from the Rhizobium species with TM 11,12, along with TM10 and ECH4 forming the dimer interface^16^. In all these structures, lipid-like densities are observed and some have been modelled providing an understanding towards the path for lipid translocation. Further, an assay for PaMprF in liposomes has shown that it can translocate a wide range of lipids^15^. Apart from the dimers, small populations of higher oligomers have been found both in Blue-Native PAGE (BN-PAGE) and in cryo-EM data sets of Pseudomonas enzyme purified in detergent as well in nanodisc^16^. The different oligomeric states observed in these studies, and the ability of the monomeric enzyme to carry out the translocation of the lipids raises the question on the in vivo relevance of the oligomeric states, substrate binding and the specificity, and the role of the flexibility of the synthase domain, observed in the cryo-EM studies.

In this study, we initially screened for different homologues of MprF that are evolutionarily closer either to Pseudomonas or Rhizobium and characterised them biochemically. Of these, the homologue of the MprF enzyme from *Agrobacterium fabrum* (strain C58), was found to be stable and amenable for structural studies. The enzyme encoded by the gene lpiA (for low pH inducible protein A and here we call it AfMprF) is involved in the synthesis of lysyl-phosphatidylglycerol (Lys-PG), which plays a vital role in bacterial infection of plants^18–20^. The MprF homologue, lpiA is found in an operon with another gene called acvB, which has been characterised as a hydrolase. Together these genes are involved in Lys-PG homeostasis, and are implicated in virulence^18,20^. The increase in Lys-PG either by overexpression of LpiA or by the absence of AcvB affects the virulence^18^. Thus, the MprF homologues from *Agrobacterium* are useful model system to study the modification of lipid and its effect on virulence. The cryo-EM structures of AfMprF reveals a dimer similar to the Rhizobium sp. and analysis of some conserved residues in the synthase domain for their ability to produce modified lipids by thin-layer chromatography (TLC) reveals a structural motif critical for substrate interaction.

## RESULTS

### Protein structure and organisation of *AfMprF*

To understand the architecture of MprF from different organisms, we expressed and characterised MprF homologues from *Azotobacter beijerinckii, Thalassovita mangrovi, Agrobacterium fabrum (formerly called A. tumefaciens* strain C58*), Enterococcus faecalis*, and *Corynebacterium glutamicum* based on their evolutionary closeness to either *Pa*MprF or *Rt*MprF (Fig. S1 A, B). Among these enzymes screened, only the MprF homologue from *A. fabrum* (*AfMprF*) was identified to be stable in detergent after solubilisation and purification (Fig. 1). The full-length *AfMprF* was obtained by heterologous expression in *E. coli*, and the lipids were extracted from the membranes overexpressing the protein as well as from *A. tumefaciens* cells. The lipid extracts resolved on a thin-layer chromatography (TLC) plate revealed peaks that stained for the amino group both from native *A. tumefaciens* and *E. coli* overexpressing AfMprF, as compared to the pET vector control, confirming that the enzyme is active and produces aminoacylated lipids (Fig. 1A)

**Figure 1:**
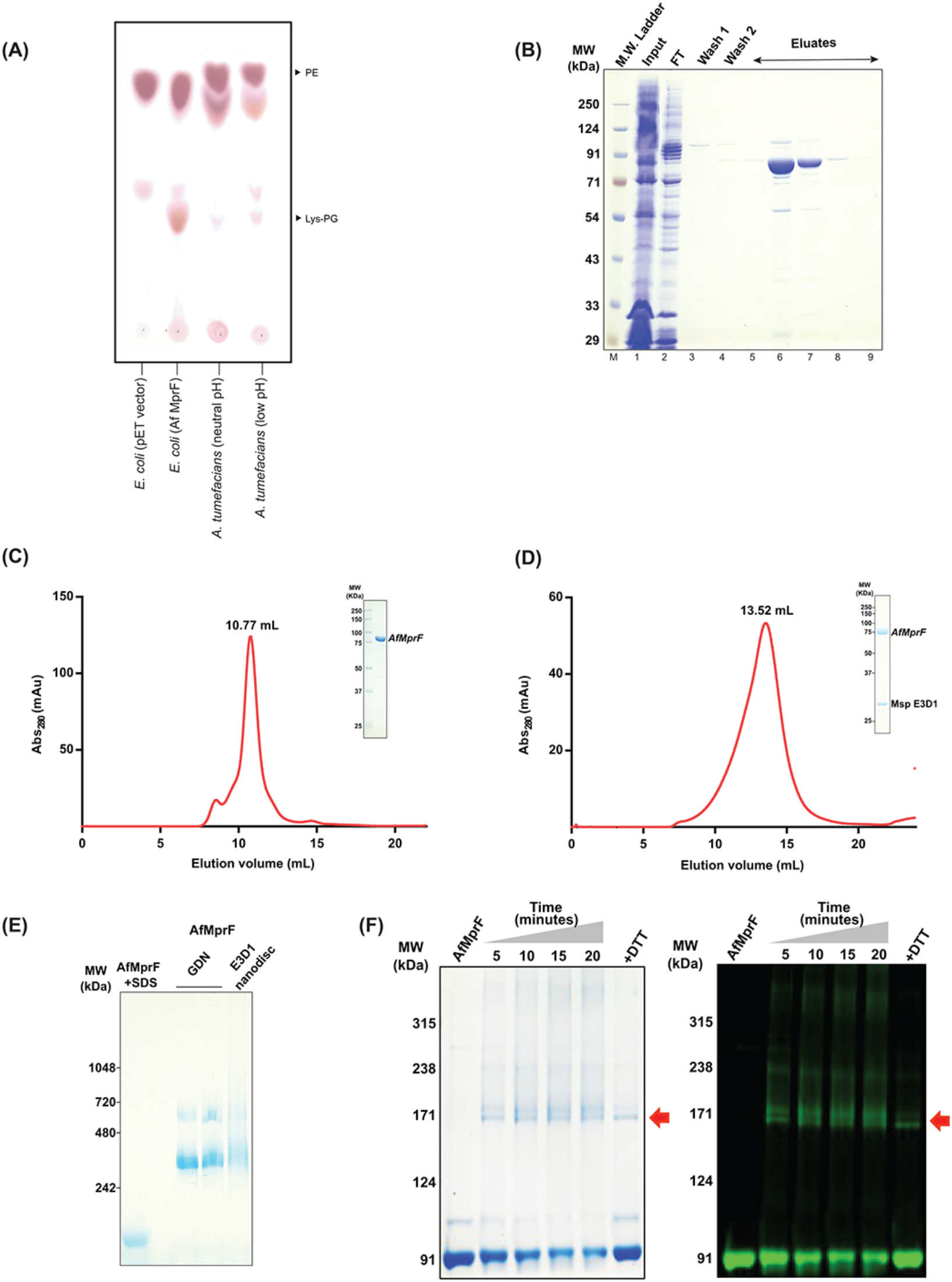
Biochemical characterisation of AfMprF. **A)** Lipid extracts from *E. coli* cells expressing *AfMprF* and vector control, along with *A. tumefaciens* lipid extracts from cells grown at two pH, resolved on TLC and stained using ninhydrin, which detects primary amine. **B)** SDS-PAGE gel image of affinity-purified *AfMprF* in GDN detergent. **C)** Size exclusion chromatogram of the *AfMprF* protein in GDN resolved on the Superdex 200 increase 10/300 column with an inset of SEC-purified *AfMprF* resolved on SDS-PAGE. **D)** Size exclusion chromatogram of the nanodisc reconstituted *AfMprF* after affinity chromatography, resolved on Superose 6 10/300 column with inset of SEC purified *AfMprF* nanodisc resolved on SDS-PAGE. **E)** The *AfMprF* protein samples in detergent and nanodisc resolved on BN-PAGE, suggesting that the protein is stable in both conditions. **F)** Cross-linking of *AfMprF*-GFP in GDN detergent using DSP resolved on a 5% SDS gel. Both the Coomassie stained and the In-gel fluorescence image of the same gel are shown. A gradual decrease in the monomeric band is observed with increase in incubation time but a complete shift of the monomeric band is not observed. The gels also show that higher order oligomers are also formed, but in the presence of DTT, the cross-linked product is dissociated and the intensity of the monomer band increases.

The *AfMprF* was extracted from the membranes using lauryl maltose neopentyl glycol (LMNG) and exchanged to glyco-diosgenin (GDN) during the metal affinity chromatography utilising the poly-histidine tag in the construct, followed by size–exclusion chromatography (SEC) (Fig. 1B and 1C). The SEC-purified protein was also reconstituted in MSP1E3D1 nanodisc using *E. coli* polar lipids, and further purified by affinity and SEC chromatography to remove empty nanodisc (Fig. 1D). The stability of the protein in detergent and lipid nanodisc was assessed by resolving the samples in BN-PAGE, which showed higher molecular weight bands, and when dissociated with SDS, they migrated as monomers (Fig. 1E). Detergent screening of AfMprF revealed the enzyme is stable in many detergents and a similar profile as in LMNG-GDN was observed (Fig S1C). To further validate the formation of an oligomer, a cross-linking experiment was performed with a construct of AfMprF with GFP at its C-terminus. The purified protein was cross-linked with Dithiobis(succinimidylpropionate) (DSP), and higher molecular weight bands were observed, revealing that AfMprF forms oligomers in detergent micelle. Addition of DTT (which cleaves the cross-linker) resulted in accumulation of the monomers of AfMprF (Fig. 1F).

Cryo-EM analysis of both the detergent-purified and the nanodisc reconstituted enzyme was performed, and the reference-free 2D suggested that the dimer of AfMprF is similar to that of *Rhizobium tropici*^10^ (Fig S2, S3). In the current data sets and classes, we do not observe other higher-order oligomers or alternate dimers. The cryo-EM maps of AfMprF in GDN and nanodisc were refined to an overall global resolution of 3.8 Å and 3.4 Å, respectively (Fig. 2 A, 2B, Fig. S2 A-F, S3 A-F, Table 1). As observed with other MprFs, the local resolution estimation of the cryo-EM maps in both detergent and nanodisc suggests that the resolution is higher in the core TM region along the dimer interface but lower towards the periphery of the membrane domain and the soluble domain (Fig. S2 G, S3 G) ^10,15,16^. The fit of the dimeric AfMprF model in cryo-EM maps is shown in Figs S2-H and S3-H. In the soluble domain, no additional densities for tRNA or lipid molecules are observed in both the maps but there are non-protein densities in the TM region, some of which could be lipids but they are currently unmodelled (Fig S2-I, S3-I). The models of AfMprF derived from GDN micelle and in nanodisc are similar with an overall RMSD of 0.3 Å. Due to higher resolution, the model derived from nanodisc has been used for analysis with constant cross-check with the map/model from GDN to locate any differences.

**Table 1:**
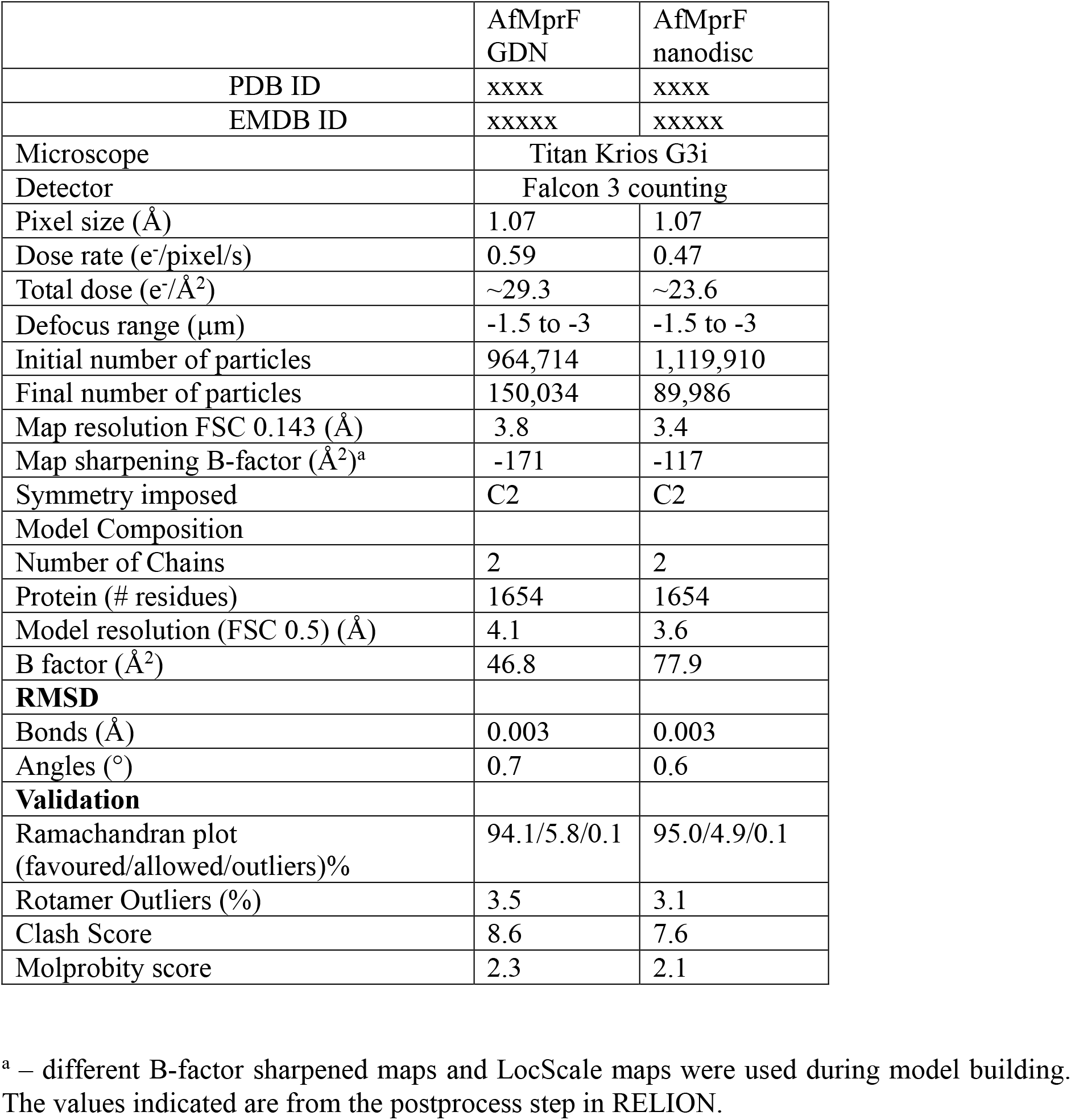
Data collection and Model refinement Statistics.

**Figure 2:**
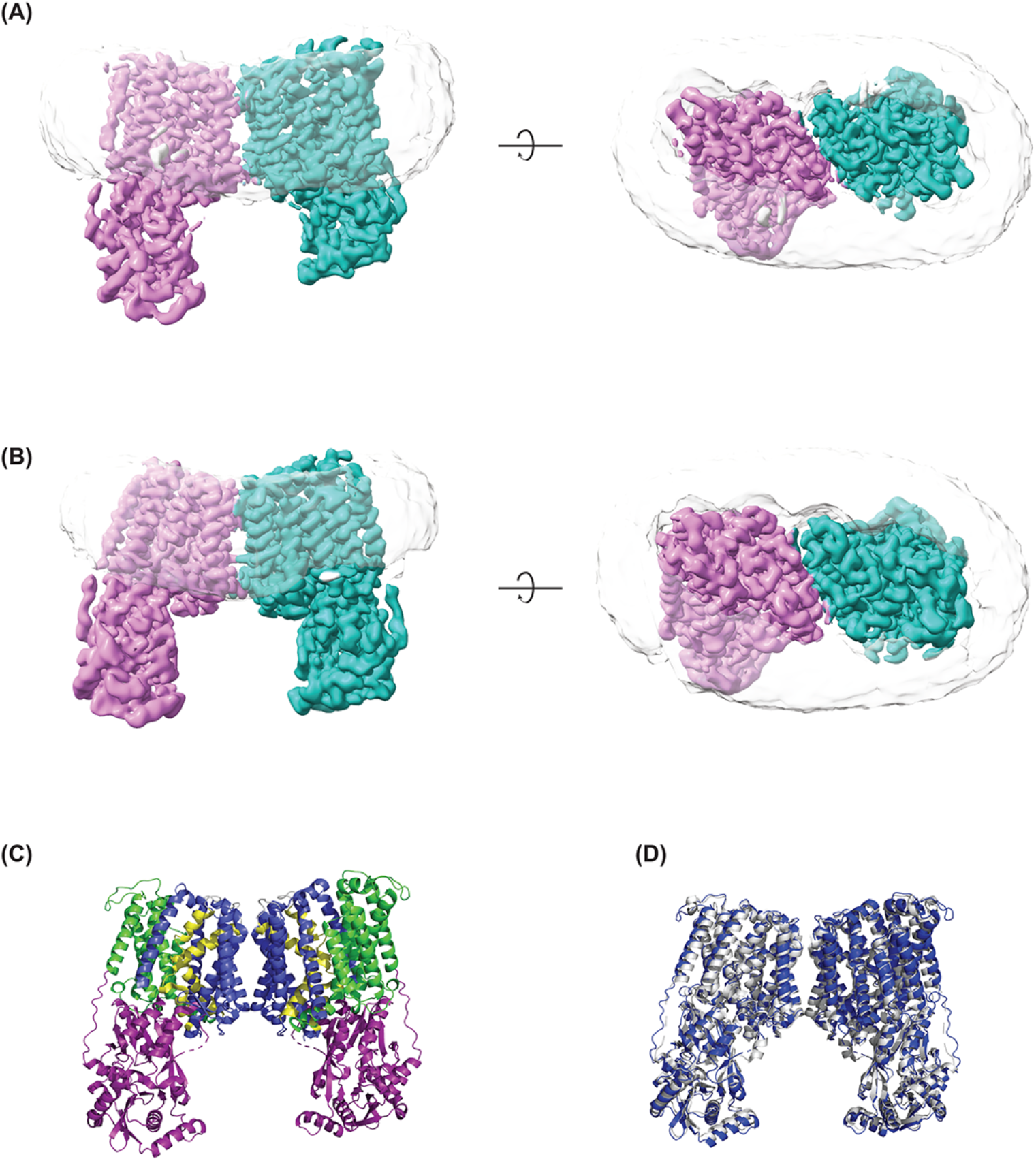
Cryo-EM maps of AfMprF. 3D reconstructions of AfMprF were performed both in GDN micelles and in nanodisc. **(A)** and **(B)** Two views of the cryo-EM maps of GDN and nanodisc are shown, respectively. Each monomer of the map is coloured in violet and sea green, the detergent/lipid belt in both maps are shown in grey with 80% transparency. The maps were generated with Locscale 2.0 and Locscale-SURFER in ChimeraX to show the enzyme and the detergent/lipid belt. **C)** The model of *AfMprF* dimer, where the N-terminal and C-terminal region of the transmembrane domain is coloured in blue and green respectively, and the re-entrant helices in yellow. The soluble synthase domain is coloured in purple. **D)** An overlay of the *AfMprF* in blue with *Rt*MprF in grey (PDB – 7DUW) showing similar protein architecture and dimer interface with an overall RMSD value of ~1.4 Å.

The structure of AfMprF reveals the typical architecture of MprF with distinct domains (Fig. 2C and S4). The dimer interface is mediated by TM helices 4, 5, 6 and 7a, similar to the *Rhizobium sp*. with few unmodelled lipid-like densities near the dimer interface (Fig 2C). Comparison of the *AfMprF* and *Rt*MprF indicates a few differences in the soluble domain with an overall RMSD of 1.4 Å (Fig 2D). Greater differences are observed when *AfMprF* and *Pa*MprF monomers are compared with an RMSD of ~4.2 Å (Fig. S5 A). The major differences between these two structures are seen in the soluble GNAT2 domain and the TM helices mediating the dimer interface. A comparison of the FL AfMprF model with the crystal structure of the soluble domain (PDB-5VRV) reveals no major changes with a RMSD of 0.73 Å (Fig, S5 B). Similarly, an overlay with the synthase domain of *B. licheniformis*^8^ reveals a very good superposition with a RMSD of 1.1 Å, and differences in the mobile elements in the GNAT2 domain (Fig. S5 C). Analysis of cavities in AfMprF reveals two large cavities in the TM domain and also in the synthase domain, where lysinamide was observed in the crystal structure (Fig. S5 D, E). Though, these cavities are not connected in the current models, it is possible that some restructuring of the helices in the GNAT2 domain can result in a continuous path between the synthase and the TM domains (Fig. S5 D).

### Conserved residues critical for aminoacylation activity in *AfMprF*

Sequence analysis of different MprF homologues shows that the synthase domain harbours several conserved residues (Fig. 3A). This is substantiated by the crystal structure of the *B. licheniformis* MprF (*Bl*MprF) soluble domain bound to the lysinamide, which suggests that the cleft in the GNAT2 domain plays an important role in the aminoacylation activity of the protein^8^. In the *AfMprF* model, the ligand is likely to be in proximity to the Asp750 and Arg753 residues (derived by overlay of the *Bl*MprF synthase domain with lysinamide) and may play a crucial role in catalysis (Fig. 3B). Mutation of these residues in *AfMprF* to alanine (D750A and R753A) results in loss of formation of LysPG (Fig. 3A, D). Further, Met752, which lies close to the lysinamide-bound cleft, does not seem to interact with the bound ligand but forms an interaction with Phe767 of the neighbouring helix of the GNAT2 domain (Fig. 3C). This interaction is termed as sulphur-aromatic motif, commonly found in certain protein families^21,22^. The mutation of this methionine residue to leucine (M752L) leads to a loss of activity as analysed by the TLC assay, suggesting that the methionine is important for the enzymatic activity (Fig. 3D).

**Figure 3:**
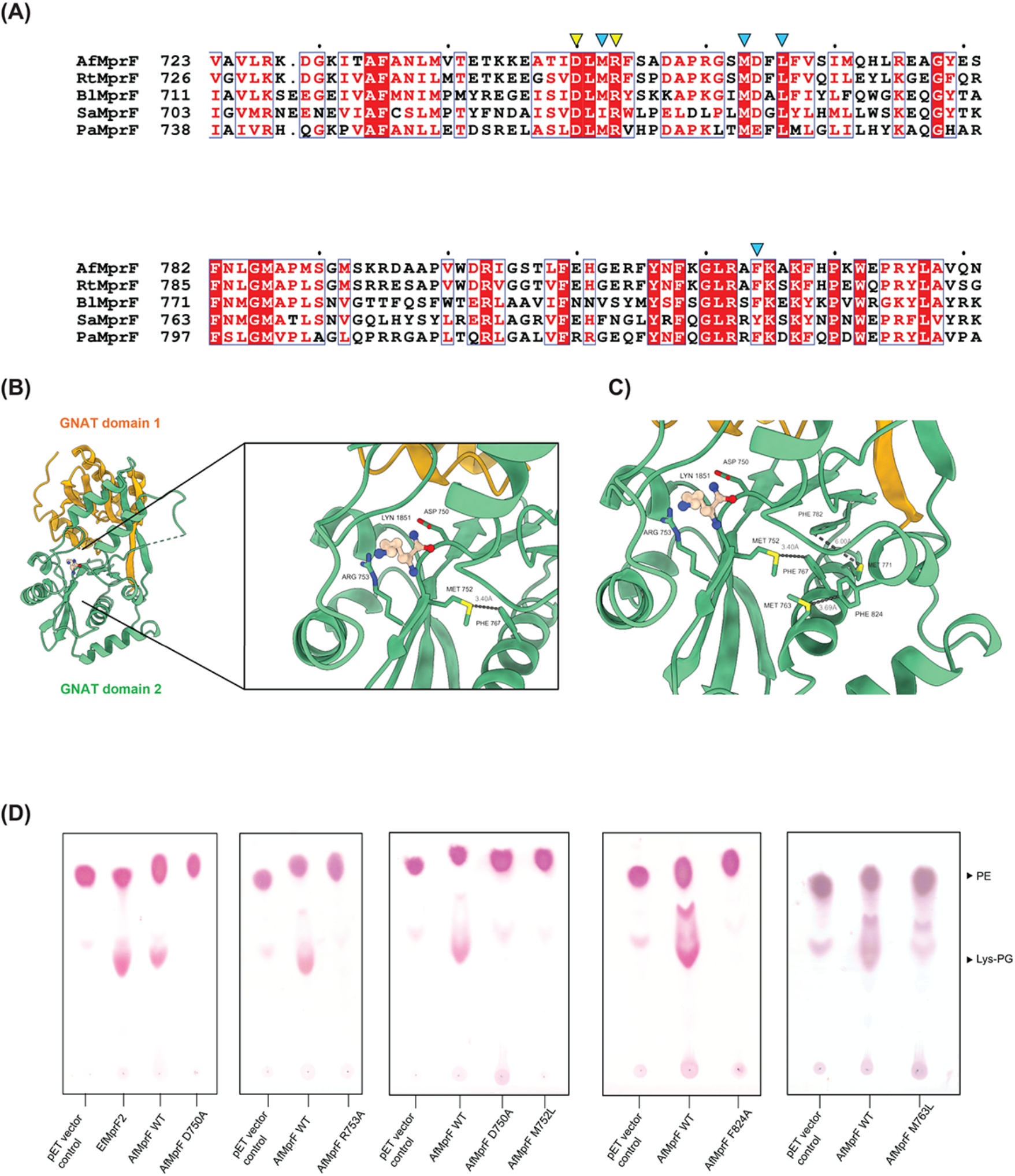
**A)** Multiple sequence alignment of select MprF homologs performed using Clustal Omega and visualised using ESPript. The conserved aspartate and arginine residues near the lysinamide binding pocket are indicated in yellow, while the residues forming the sulphur–aromatic motifs are indicated in blue. **B)** Overlay of the soluble domains of *AfMprF* with PDB – 4V36 showing only the lysinamide molecule. The GNAT domains 1 and 2 of *AfMprF* are shown in orange and green, respectively. The lysinamide binding cleft in the GNAT domain 2 of *AfMprF* shows Asp750 and Arg753 in the ligand proximity, while Met752 is away from the binding site and interacts with the Phe767 from the neighbouring helix. **C)** Possible sulphur-aromatic motif pairs near the ligand binding site mediated by Met752 – Phe767, Met771 – Phe782, and Met763 – Phe824 are shown. **D)** TLC analysis of the lipids extracted from *E. coli* cells overexpressing *AfMprF*-WT, *AfMprF* D750A, *AfMprF* R753A, *AfMprF* M752L, *AfMprF* F824A, and M763L, along with the vector as a negative control and *Ef*MprF2 as a positive control.

The sulphur aromatic residues have been extensively characterised and are important in stabilising protein structure^21,22^. Analysis of the soluble domain of the *AfMprF* model reveals the presence of three sulphur-aromatic motifs near the proposed tRNA binding site in the GNAT2 sub-domain. Apart from the Met752 and F767 pair described above, the following pairs Met771/Phe782, and Met763 and Phe824 also form this motif. Of these, Met763/Phe824 is highly conserved in MprF homologs but Met771/Phe782 is present only in *Agrobacterium* MprF (Fig. 3A and 3C). Phe824 has previously been hypothesized to form a ribose-π interaction with the tRNA substrate in the cleft of the GNAT2 domain^8^. In *AfMprF*, the mutation of this phenylalanine residue to alanine (F824A) leads to a loss in the activity, suggesting that this residue is critical for the enzymatic function (Fig. 3D). Similarly, mutation of Met763 to leucine leads to a loss of activity. Thus, we propose that the sulphur aromatic motifs in the protein active site are important to maintain protein tertiary structure as well as for protein substrate interaction, particularly with the incoming aminoacyl-tRNA substrate.

The transfer of the amino acid onto the lipid head group requires the lipid to be in the proximity of the active site. Previous studies and our analysis of conserved residues in the GNAT2 domain suggest the possible binding site for the tRNA substrate^8,9^. However, the exact mechanism by which the amino acid is transferred from the tRNA substrate and the lipid binding site is unclear. The *AfMprF* model reveals that the distance between the proposed active site in the GNAT2 domain to the lipid bilayer is around ~43 Å, suggesting that the enzyme needs to undergo some subtle or large changes to couple the transfer of amino acid from the cytosol tRNA pool onto the lipid molecule. Previous studies on the synthase domain of *Bl*MprF using molecular docking suggest that the cavity in GNAT1 domain can accommodate a lipid molecule and helps the protein to transfer the amino acid directly onto the lipid molecule^8^. In the GNAT1 domain, a conserved hydrophobic residue, Leu645 is observed at the tunnel region, where lipid molecules can bind (Fig. S1A). Mutation of the Leu645 residue at the opening of this tunnel to arginine leads to a loss of protein activity (Fig. 4A, B). However, mutation of Leu645 to hydrophobic side chains (L645I, L645F) or negatively charged amino acid (L645E) resulted in the formation of lys-PG (Fig. 4b). This suggests that positive charge is detrimental at this position, and the loss of activity could be because of lack of substrate binding or change in the local protein structure.

**Figure 4:**
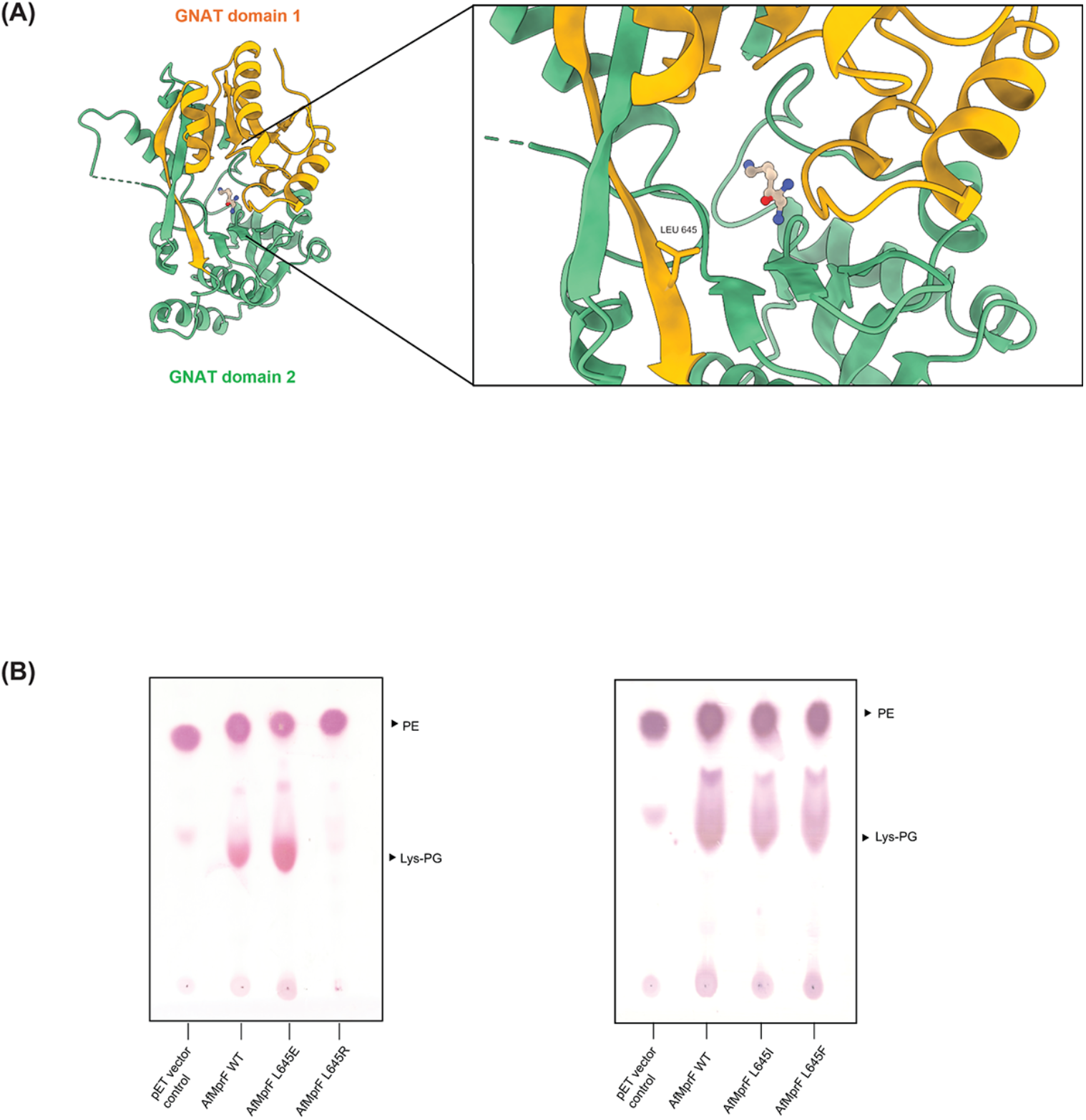
Analysis of a key residue on the GNAT1 domain. **A)** The soluble domain of the *AfMprF* with Leu645 near the cleft of GNAT domain 1 is shown. **B)** TLC analysis of the lipids extracted from *E. coli* cells overexpressing *AfMprF*-WT, *AfMprF* L645E, *AfMprF* L645R, *AfMprF* L645I, and *AfMprF* L645F, along with the vector as the negative control. This analysis shows that hydrophobic residues are tolerated at this position and also negative charge but not a positive charged residue that results in no production of Lys-PG.

## Discussion

The oligomeric state of a protein plays an important role in protein stability, function, and also helps in cooperative interaction and allosteric regulation^23–25^. Previous work on MprF from *Pseudomonas and Rhizobium* suggests that the enzymes are dimeric but have different dimer interfaces^10,15,16^. To understand the role of the oligomers in the protein function, we expressed and purified six MprF homologs with different sequence similarity to *Rt*MprF and *Pa*MprF (Fig S1B). However, due to the low stability of many MprF proteins in detergent, we were only able to biochemically and structurally characterise *AfMprF* in vitro. The structures of the *AfMprF* and in vitro cross-linking experiment of the purified protein suggest that the protein is a dimer and adopts a similar overall architecture and dimer interface as *Rt*MprF (Fig. 1, 2).

Mutagenesis study of residues in *AfMprF* close to the lysinamide binding site suggests that apart from the aspartate and arginine residue, methionine also plays a vital role in the enzyme function by forming a sulphur aromatic motif with the phenylalanine of the neighbouring helix and stabilizing the protein active site (Fig 3). We identify an additional sulphur aromatic motif in *AfMprF* mediated by Met763 and Phe824, and a mutation in either of these residues leads to a loss in protein activity. Given that the Phe824 residue has been previously implicated in stabilizing the protein-tRNA complex, we propose that the Met763 helps stabilize the Phe824 residue conformation upon tRNA binding.

A recent report of the *Pa*MprF structure in lipid nanoparticles revealed a monomeric enzyme and the reconstituted enzyme in liposomes can perform the flippase activity and promiscouous^15^. Further, it has also be shown using cysteine cross-linking experiments that the *Pa*MprF can form a dimer in *E. coli* membranes overexpressing the enzyme^16^. It is clear that all the elements necessary for the tRNA binding and lipid flipping exist in the Mprf monomer, and the oligomer formation might increase stability or be involved in regulation. For example, the MprF homologous genes are typically found alongside a hydrolase and thereby the levels of modified lipids are controlled^18,26^. Multiple sequence analysis of the MprF homologs revealed that the *AfMprF* shares more than 77% and 56% sequence similarity with the *Rt*MprF and *Pa*MprF, respectively. Thus, *AfMprF* is closely related to the *Rt*MprF, suggesting that closely related homologues might adapt a similar oligomeric state (Fig. 2C, S1 B). However, this cannot be generalised yet as the dimer interface is not conserved and characterisation of other homologues is required. Further characterization of these MprF homologs in native vesicles or proteoliposomes will shed further light on the oligomeric state, the role of lipids in the formation of these oligomers and perhaps in trapping the substrate to advance our understanding of the mechanism of these enigmatic enzymes.

## Material and methods

### Protein expression and purification of *AfMprF*

The codon optimised MprF gene of the *A. fabrum (*formerly called *A. tumefaciens* strain C58, Uniprot ID - A9CHP8) was synthesised and cloned into a modified pET29b vector with a C-terminal strep-tag II and polyhistidine tag (Twist Bioscience). Mutations were generated using PCR-based site-directed mutagenesis, and the constructs were verified using sequencing.

The *AfMprF* constructs were overexpressed using the *E. coli* BL21(DE3) strain. For protein purification, the cells were inoculated in Terrific Broth containing 50 μg/ml of kanamycin and grown at 37 °C at 180 rpm. Protein expression was induced using 1mM IPTG, and the culture was grown overnight at 25 °C. All the steps were carried out on ice or at 4 °C unless otherwise specified. The cells were harvested by centrifugation at 4000 g for 30 minutes and resuspended in lysis buffer containing 50 mM Tris pH 8.0, 200mM NaCl, 4 mM MgCl_2_, 1 mM CaCl_2_, 2 μg/ml DNase I, 1mM PMSF, and 5mM ß-mercaptoethanol. Cells were lysed by mechanical disruption at 15,000 psi using Emulsiflex-c3 (Avestin). The cell debris and unbroken cells were removed by centrifugation at 15,000 g for 20 minutes. The clarified supernatant was subjected to ultracentrifugation at 100,000 g for 90 minutes to isolate the membranes. The membranes were resuspended in buffer A containing 40 mM Tris pH 8.0, 200 mM NaCl, and stored at -80 °C.

For *AfMprF* purification (all steps performed at 4 ^°^C unless otherwise stated), the membranes were resuspended in buffer A containing 10 mM imidazole at a 1:20 w/v ratio and solubilised using 1 % LMNG (Anatrace) by stirring for 1 hour. The unsoluble material was pelleted by ultracentrifugation at 100,000 g for 30 minutes, and the supernatant was applied to 2 ml of Ni-NTA agarose resin (Qiagen) preequilibrated with buffer A + 0.01% GDN for 90 minutes. The Ni-NTA slurry was then transferred to an Econo-Pac column (Biorad) and washed sequentially with 25 ml using wash buffer 1 (40 mM Tris pH 8.0, 200 mM NaCl, 20 mM imidazole, 0.01 % GDN) and wash buffer 2 (40 mM Tris pH 8.0, 200 mM NaCl, 40 mM imidazole, 0.006 % GDN). *AfMprF* was eluted with 10 ml of elution buffer containing 40 mM Tris pH 8.0, 200 mM NaCl, 0.006 % GDN, and 250 mM Imidazole. The protein-containing fractions were pooled and concentrated using a 100-KDa MW cutoff concentrator (Millipore). The protein was further purified using SEC and injected onto a Superdex 200 increase 10/300 column (Cytiva) pre-equilibrated with SEC buffer (20 mM Tris pH 8.0, 150 mM NaCl) containing 0.005 % GDN. The peak fractions were pooled and concentrated to around 5.7 mg/ml using a 100-KDa MW cut-off concentrator for cryo-EM grid preparation.

### Cross-linking of *AfMprF*

The *AfMprF* cDNA was cloned into a pWARF(-) vector with a C-terminal HRV3C protease site, followed by GFP and polyhistidine tag^27^. After SEC purification, the buffer was exchanged to 20 mM HEPES, pH 8.0, 150 mM NaCl, and 0.005 % GDN. The protein was chemically crosslinked using 40 μM Dithiobis(succinimidylpropionate) (DSP) and incubated at room temperature. The reaction was stopped using 100 mM Tris, pH 7.4, and mixed with 6X non-reducing SDS-PAGE buffer. The disulphide bond of DSP was cleaved by the addition of 50 mM DTT, followed by incubation at 37 ^°^C for 30 minutes. All the samples were resolved on a 5 % SDS gel and imaged for in-gel fluorescence and stained using Coomassie stain.

### MSP purification

MSP1E3D1 (MSP) containing an N-terminal His tag was expressed in BL21 (DE3) cells and was purified using Ni-NTA chromatography according to previously published protocol ^28,29^. The polyhistidine tag was cleaved by TEV protease, and the protein was further purified by reverse IMAC. The cleaved MSP protein was dialyzed against a buffer containing 20 mM Tris pH 7.4, 200mM NaCl, and concentrated to 10mg/ml. The protein was flash-frozen in liquid nitrogen and stored at -80°C for further use.

### *AfMprF* reconstitution in lipid nanodisc

*E. coli* polar lipids (EPL) were resuspended in SEC buffer containing 2 % CHAPS and mixed with MSP1E3D1 at room temperature for 10 minutes. SEC-purified *AfMprF* in GDN was added to the lipid mixture and incubated for 30 minutes at 4 °C. Nanodisc reconstitution was initiated by the addition of 200 mg of Bio-beads (Bio-Rad) pre-equilibrated with SEC buffer, and the mixture was rotated for 30 minutes at 4 °C. An additional 200mg of beads was added, and the mixture was incubated overnight at 4 °C. The mixture was centrifuged, and the protein was applied to 1 ml of Ni-NTA resin pre-equilibrated with equilibration buffer (20 mM Tris pH 8.0, 150 mM NaCl, 10mM imidazole) and incubated for 60 minutes. The resin was washed with 20 column volumes of wash buffer (20 mM Tris pH 8.0, 150 mM NaCl, 20 mM imidazole) and the protein was eluted with elution buffer (20mM Tris pH 8.0, 150 mM NaCl, 250 mM imidazole). The eluates were pooled, concentrated, and injected into the Superose 6 increase column preequilibrated with SEC buffer. The peak fractions were concentrated to around 6.3 mg/ml using a 100-KDa MW cut-off concentrator for cryo-EM sample preparation.

### Cryo-EM sample preparation and data acquisition

The *AfMprF* protein purified in GDN or reconstituted in nanodisc was centrifuged at 25,000 g for 10 minutes. A holey carbon, 300 mesh R1.2/1.3 gold grid (Quantifoil) was glow-discharged for 60 seconds (PELCO), and 3 µl of protein sample was applied to the grid at 16 °C and 100% humidity. The grids were blotted for 3-4 seconds and plunge-frozen in liquid ethane using a Vitrobot Mark IV (ThermoFisher Scientific). Grids were stored in liquid nitrogen until imaging. On the day of screening and imaging, grids were mounted on the Autogrids and transferred to Titan Krios (ThermoFisher Scientific). Grids were screened using Low dose protocol, and data collection of selected squares with good ice and particle distribution was set up with EPU (ThermoFisher Scientific). Imaging was performed with Falcon 3 detector in counting mode at 75,000 x corresponding to a pixel size of 1.07 Å for 60 seconds, movie frames were saved for further processing.

### Cryo-EM data processing, Model building and analysis

For the *AfMprF* GDN dataset, a total of 1778 movies were collected and imported into RELION ^30^. Motion correction was performed using Relion, followed by contrast transfer function estimation using Gctf ^31–34^. Micrographs with resolution higher than 4 Å, and defocus values between 1.0 – 3.6 µm were selected, resulting in 1718 micrographs. Laplacian-of-Gaussian (LoG) based autopicking was performed on a set of 250 micrographs, and the autopicked particles were extracted and subjected to 2D classification. Final 16,486 particles were used as templates for reference-based autopicking, and 964,714 particles were autopicked and extracted with a box size of 300 pixels, followed by iterative 2D classification to give 392,995 particles. These particles were imported into cryoSPARC^35,36^ and subjected to two rounds of ab-initio reconstructions and heterogeneous refinements (C1 symmetry), followed by non-uniform refinement with C2 symmetry, resulting in a map with an overall resolution of 3.99 Å (150,034 particles). This map was post-processed in RELION using a mask, and the output of which was used for Bayesian polishing^37^. The particles were then refined in cryoSPARC using local and global CTF refinements and non-uniform refinement to yield a map with 3.8 Å overall resolution.

For the *AfMprF* nanodisc dataset, a total of 2067 movies were collected and imported into RELION. Motion correction was performed using RELON, followed by contrast transfer function estimation using Gctf. Micrographs with a resolution higher than 4 Å were selected, resulting in 2066 micrographs. Laplacian-of-Gaussian (LoG) based autopicking was performed on a set of 218 micrographs, and the autopicked particles were extracted and subjected to 2D classification. Final 20,827 particles were used as templates for reference-based autopicking, resulting in 1,119,910 particles. The particles were binned 2x and extracted with a box size of 320 pixels, followed by iterative 2D classification to give 392,995 particles. The particles were reextracted with a pixel size of 1.07 Å, imported into cryoSPARC and subjected to ab-initio reconstruction and heterogeneous refinement (three classes, C1 symmetry), followed by non-uniform refinement with C2 symmetry, resulting in a map with an overall resolution of 3.51 Å (185,870 particles). Second round of ab-initio model generation and heterogeneous refinement (two classes, C2 symmetry) followed by CTF refinement and non-uniform refinement with C2 symmetry yielded a map with overall resolution of 3.42 Å (115,090 particles). This map was post-processed in RELION using a mask, and the output of which was used for Bayesian polishing. The particles were then refined in cryoSPARC using local and global CTF refinements and non-uniform refinement (C2 symmetry) and a single 2D classification to yield a map with an overall resolution of ~3.4 Å from 89,986 particles.

The model of AfMprF was built using the AlphaFold^38^ model by first docking to the cryoEM map using Chimera^39^ and manual inspection in Coot^40^. The soluble domain in particular required careful inspection and rebuild. Multiple post-processed maps were used during model building including different B-factor sharpening and the map from LocScale 2.0^41^. The model was then refined in Phenix and Servalcat-Refmac^42–45^. Cavities in AfMprF were identified within ChimeraX using KVFinder^46^. Sequence analysis was performed with Clustal omega _47,48_, visualisation with ESPript^49^ and the phylogenetic tree was calculated with Mega 4.0^50^. Figures were made with ChimeraX^51^ and PyMoL^52^.

### Culturing of *Agrobacterium tumefaciens*

A single colony of *A. tumefaciens* strain GV3101 was inoculated into the primary culture and grown in containing yeast extract beef extract (YEB), and incubated at 30 ºC, 180 rpm overnight. The secondary culture media were buffered at pH 4.5 and pH 7.0 using 10 mM sodium acetate-acetic acid and HEPES-NaOH, respectively. The primary culture was used to inoculate secondary cultures at different pH levels and further incubated at 30 ºC, 180 rpm overnight. The cultures were centrifuged at 4000 rpm for 30 minutes to harvest the cells and stored at -80 ºC.

### Lipid extraction

Lipids were extracted from cells using a modified Bligh and Dyer method according to the previously described protocols^53,54^. Briefly, BL21 (DE3) *E. coli* cells expressing the wild-type or mutant protein were pelleted at 4,000 g for 10 minutes. Cells were resuspended into 200 μL of 120 mM sodium acetate, pH 4.5, and mixed with 750 μL of chloroform:methanol (1:2, v/v). The mixture was vortexed for 5 minutes at room temperature, and 250 μL of chloroform was added and vortexed again. Finally, 250 μL of 120 mM sodium acetate, pH 4.5, was added to the mixture and mixed by vortexing. The sample was centrifuged at 1000 g for 2 minutes, and the lower organic phase containing lipids was separated from and dried under a stream of argon gas. The lipids were resuspended in 30 μL of chloroform and used for the TLC assay. A similar protocol was used for the extraction of lipids from *A. tumefacians*.

### TLC assay

Lipid extracts were spotted on TLC silica gel 60 F_254_ plates (Sigma Aldrich), and the lipid extracts were resolved using a mobile phase of chloroform:methanol:water at 65:25:5 ratio. The TLC plates were stained and developed using ninhydrin (0.3% in n-butanol +1.5% acetic acid v/v).

## Acknowledgements

We acknowledge the National Cryo-EM facility, Bangalore, for data collection, which was supported by the Department of Biotechnology, DBT/PR12422/MED/31/287/2014, and the computing facility in the Bangalore Life Science Cluster, in particular Mr Chakrapani. SD acknowledges the graduate fellowship from TIFR/NCBS. PS is a recipient of ANRF-NPDF fellowship. K.R.V. acknowledges the support of the Department of Atomic Energy, Government of India, Project Identification No. RTI 4006. KRV is part of the EMBO Global Investigator Network.

## Data Availability

The cryo-EM maps and the coordinates will be deposited in the EMDB and PDB respectively with the following accession codes.

AfMprF in GDN – EMD-xxxxx and PDB xxxx and AfMprF in nanodisc – EMD-xxxxx and PDB xxxx

