## Supplementary 1 for "Biochemical and structural characterisation of MprF homologue, LpiA from *Agrobacterium*"

**(A)**

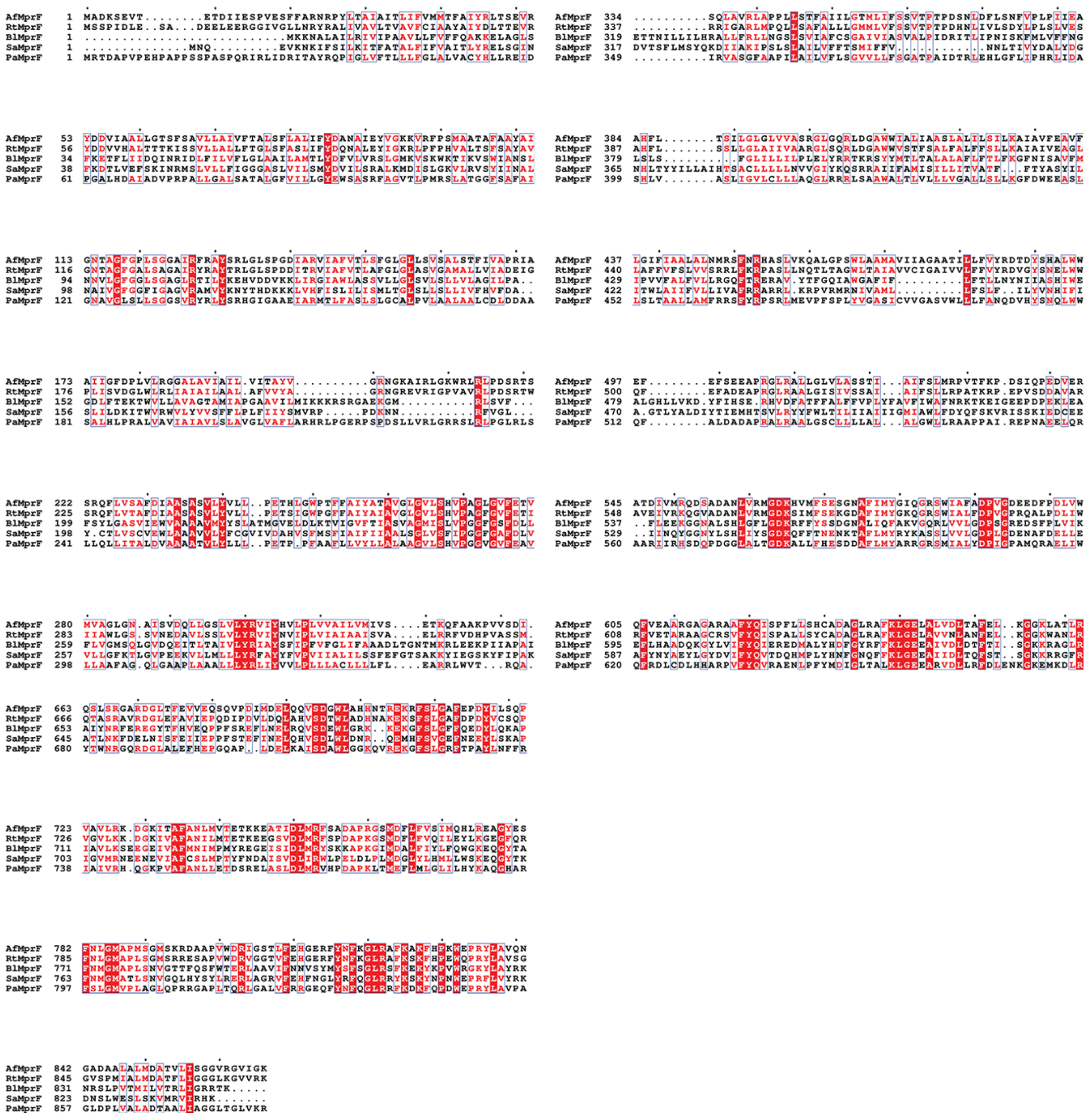

(B)

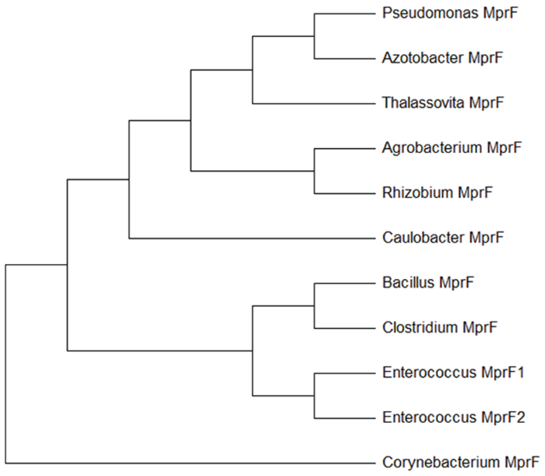


(C)

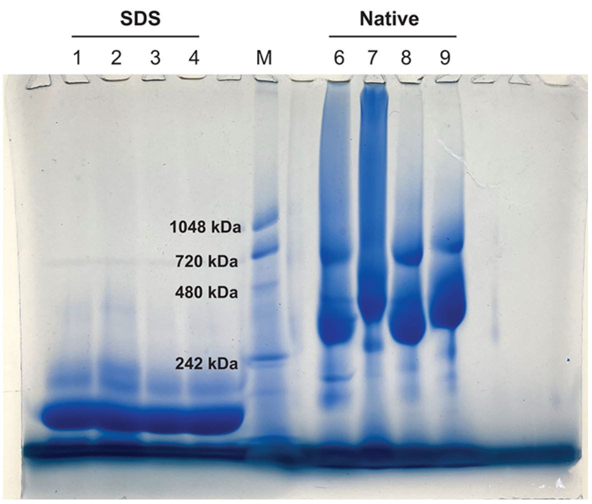


**Figure S1: A)** Multiple sequence alignment of MprF homologs was performed using Clustal Omega[47,48] and visualised using ESPript[49]. Few of these homologs were used for expression and biochemical screening. **B)** Neighbour-joining tree calculated from the alignment in panel A using Mega 4.0[50]. (C) A BN-PAGE gel showing the profile of *Af*MprF extracted and purified in different detergents. Lanes 1-4 are of samples treated with 1% SDS before loading. Lanes 6-9 are samples of AfMprF in different detergents. Lane M is marker, mitochondrial membrane proteins solubilised in DDM as the ladder. Lanes 1, 6 – solubilised and purified in DDM, Lanes 2, 7 - solubilised in DDM and purified in GDN, Lanes 3, 8 - solubilised and purified in LMNG, Lanes 4, 9 - solubilised in LMNG and purified in GDN. In all the native conditions, bands are observed above the 242 kDa marker, indicating that *Af*MprF exists as a higher oligomer in different detergents.

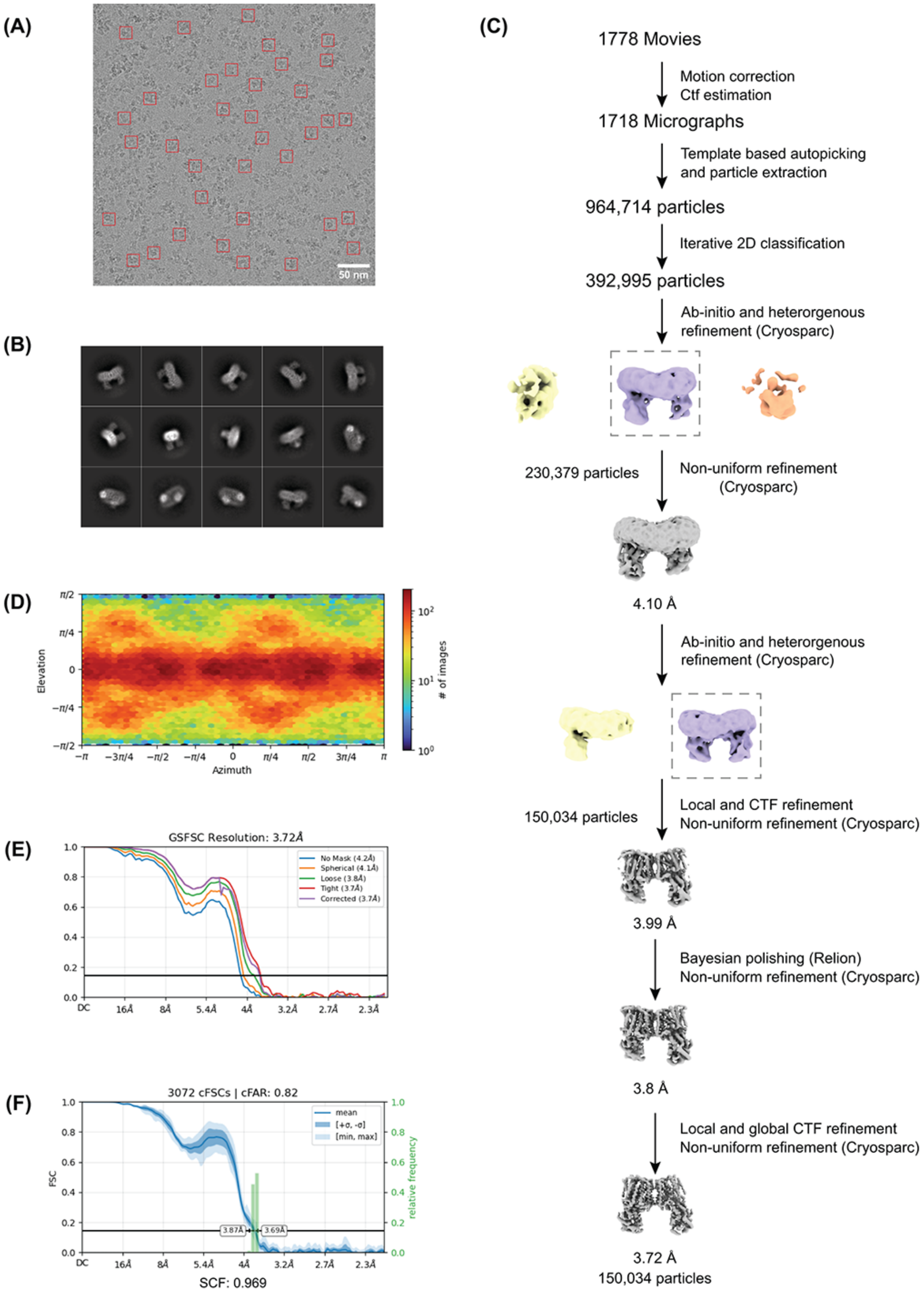
(A)
50 nm
(B)
(C)
1778 Movies
Motion correction
Ctf estimation
1718 Micrographs
Template based autopicking
and particle extraction
964,714 particles
Iterative 2D classification
392,995 particles
Ab-initio and heterorgenous
refinement (Cryosparc)
230,379 particles
Non-uniform refinement
(Cryosparc)
4.10 Å
Ab-initio and heterorgenous
refinement (Cryosparc)
150,034 particles
Local and CTF refinement
Non-uniform refinement (Cryosparc)
3.99 Å
Bayesian polishing (Relion)
Non-uniform refinement (Cryosparc)
3.8 Å
Local and global CTF refinement
Non-uniform refinement (Cryosparc)
3.72 Å
150,034 particles
(D)
Elevation
Azimuth
### of images
(E)
GSFSC Resolution: 3.72Å
No Mask (4.2Å)
Spherical (4.1Å)
Loose (3.8Å)
Tight (3.7Å)
Corrected (3.7Å)
(F)
3072 cFSCs | cFAR: 0.82
mean
FSC
relative frequency
3.87Å
3.69Å
SCF: 0.969

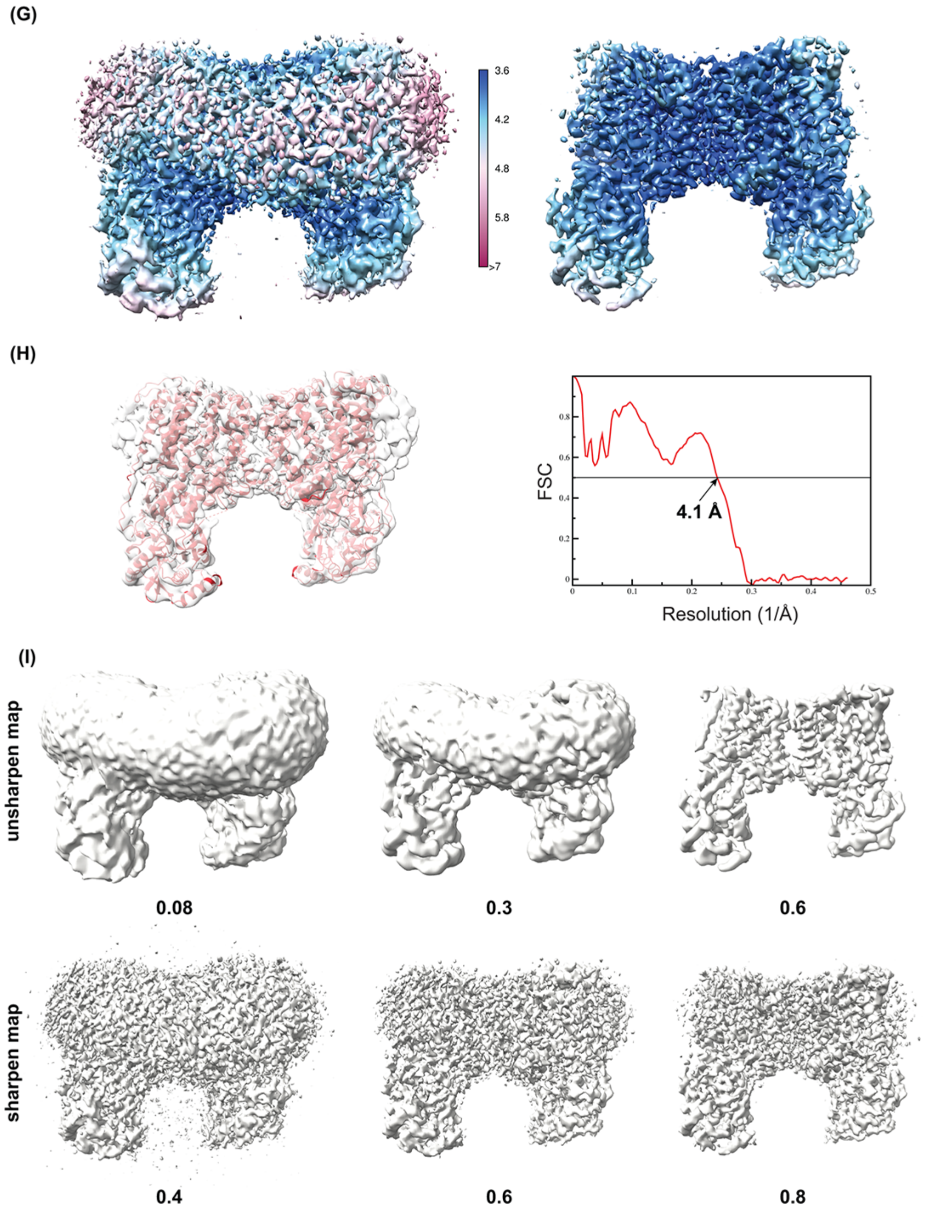


**Figure S2: Cryo-EM data processing and analysis of *Af*MprF in GDN micelles. A)** Representative micrograph of *Af*MprF enzyme in GDN at 75000x magnification with particles in red box (Scale bar = 50 nm). **B)** 2D class averages of the protein showing different orientations of the particles. **C)** Data processing workflow of the *Af*MprF in GDN using both RELION and CryoSPARC. **D)** Orientation plot of the *AfMprF* in GDN represented as a heatmap. **E)** Gold-standard FSC of the final map from a subset of particles indicates that the resolution is around 3.8 Å (3.7 Å with a tight mask) using the 0.143 criterion. **F)** The anisotropy or preferred orientation distribution was analysed using the orientation diagnosis routine in cryoSPARC, which shows a sampling compensation factor (SCF) value of 0.969 and cFAR value of 0.82. **G)** Local resolution plot of *Af*MprF calculated using RELION showing the core of the enzyme at higher resolution and the periphery, the detergent/lipid belt at lower resolution.

The maps are shown at two threshold values. **H)** The fit of the model (red) on the map (grey transparent surface) and the FSC of the map vs model at 0.5 gives a resolution of 4.1 Å. **I)** The final unsharpened and sharpened maps are shown at different thresholds (values given beneath each map). In the lower threshold maps, large flexibility of the soluble domain or other additional densities are not observed.

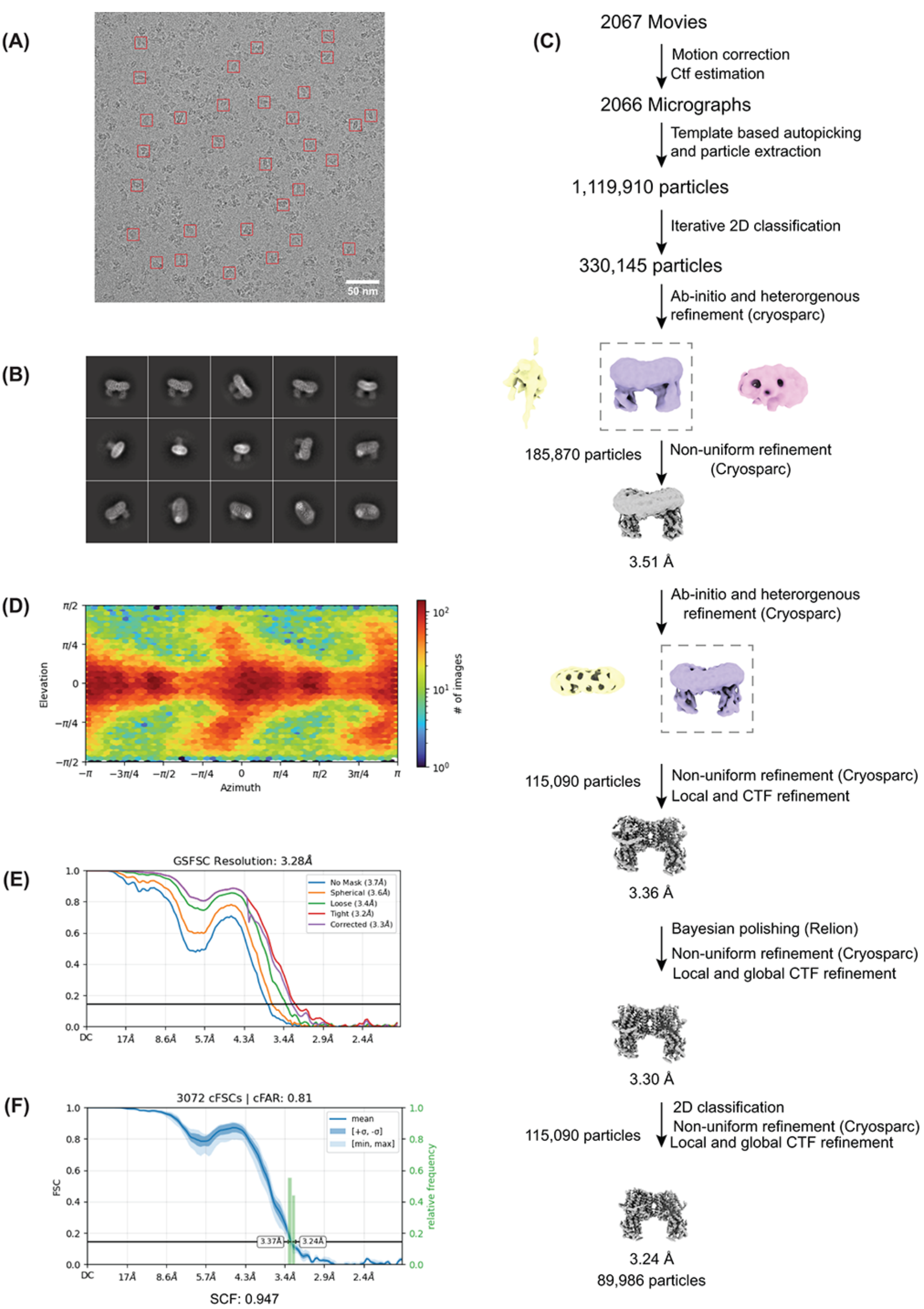
(A)
50 nm
(B)
(C)
2067 Movies
Motion correction
Ctf estimation
2066 Micrographs
Template based autopicking
and particle extraction
1,119,910 particles
Iterative 2D classification
330,145 particles
Ab-initio and heterorgenous
refinement (cryosparc)
185,870 particles
Non-uniform refinement
(Cryosparc)
3.51 Å
Ab-initio and heterorgenous
refinement (Cryosparc)
115,090 particles
Non-uniform refinement (Cryosparc)
Local and CTF refinement
3.36 Å
Bayesian polishing (Relion)
Non-uniform refinement (Cryosparc)
Local and global CTF refinement
3.30 Å
2D classification
115,090 particles
Non-uniform refinement (Cryosparc)
Local and global CTF refinement
3.24 Å
89,986 particles
(D)
Elevation
Azimuth
### of images
(E)
GSFSC Resolution: 3.28Å
No Mask (3.7Å)
Spherical (3.6Å)
Loose (3.4Å)
Tight (3.2Å)
Corrected (3.3Å)
(F)
3072 cFSCs | cFAR: 0.81
FSC
relative frequency
mean
[+σ, -σ]
[min, max]
3.37Å
3.24Å
SCF: 0.947

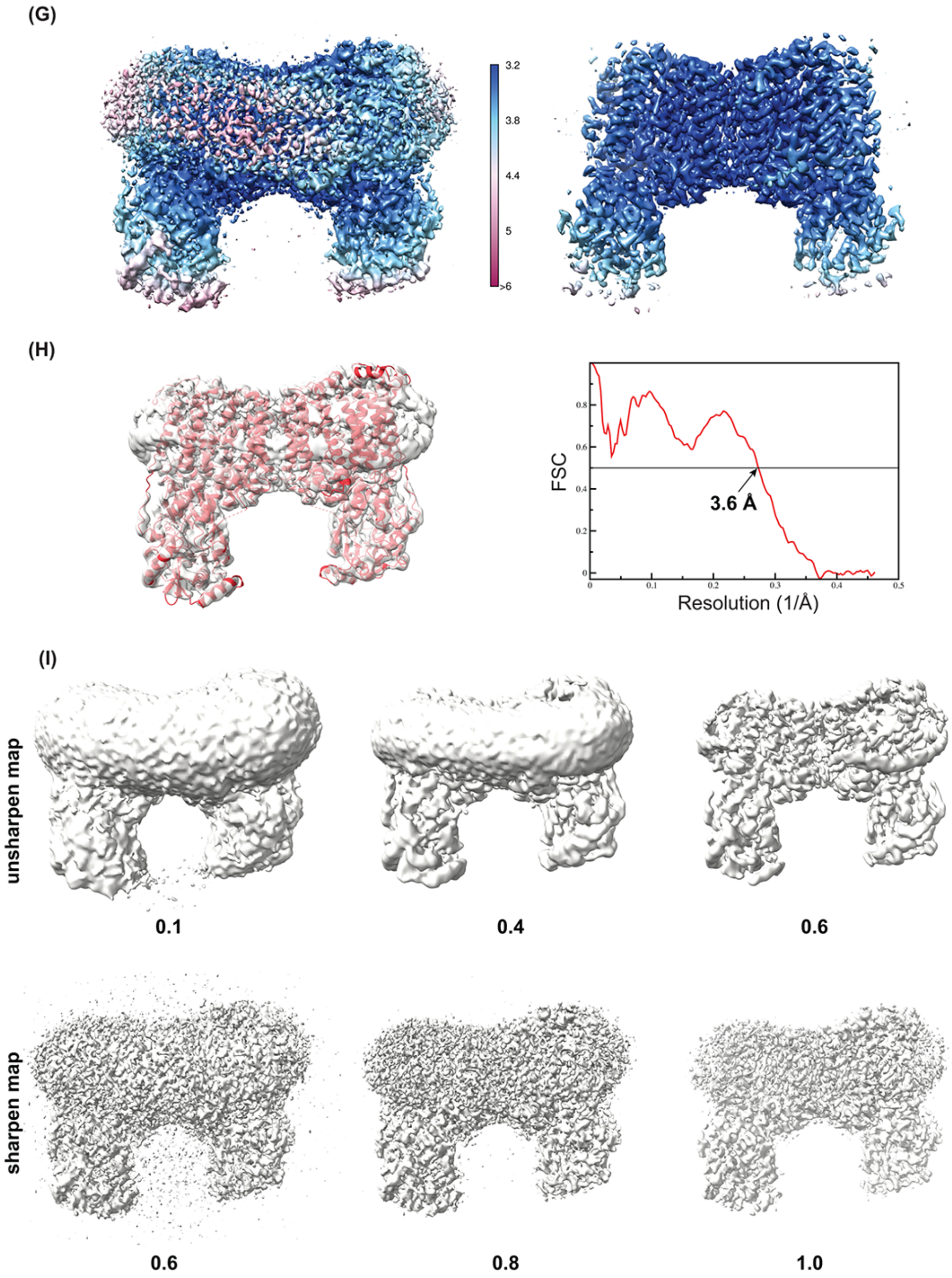


**Figure S3: Cryo-EM data processing and analysis of *Af*MprF in nanodisc. A)** Representative micrograph of *AfMprF* enzyme in nanodisc at 75000x magnification with particles in red box (Scale bar = 50nm). **B)** 2D class averages of the protein showing different orientations of the particles. **C)** Data processing workflow of the *Af*MprF in nanodisc using both RELION and CryoSPARC. **D)** Orientation plot of the *AfMprF* in nanodisc represented as a heatmap. **E)** Gold-standard FSC of the final map from a subset of particles indicates that the resolution is around 3.4 Å (3.3 Å with a tight mask) using the 0.143 criterion. **F)** The anisotropy or preferred orientation distribution was analysed using the orientation diagnosis routine in cryoSPARC, which shows a sampling compensation factor (SCF) value of 0.95 and cFAR value of 0.81. **G)** Local resolution plot of *Af*MprF in nanodisc calculated using RELION showing the core of enzyme at higher resolution and the periphery, the detergent/lipid belt at lower resolution. The maps at two threshold values are shown. **H)** The fit of the model (red) on the map (grey transparent surface) and the FSC of the map vs model at 0.5 gives a resolution of 3.6 Å. **I)** The final unsharpened and sharpened maps are shown at different thresholds (values given beneath each map). In the lower threshold maps, large flexibility of the soluble domain or other additional densities are not observed.

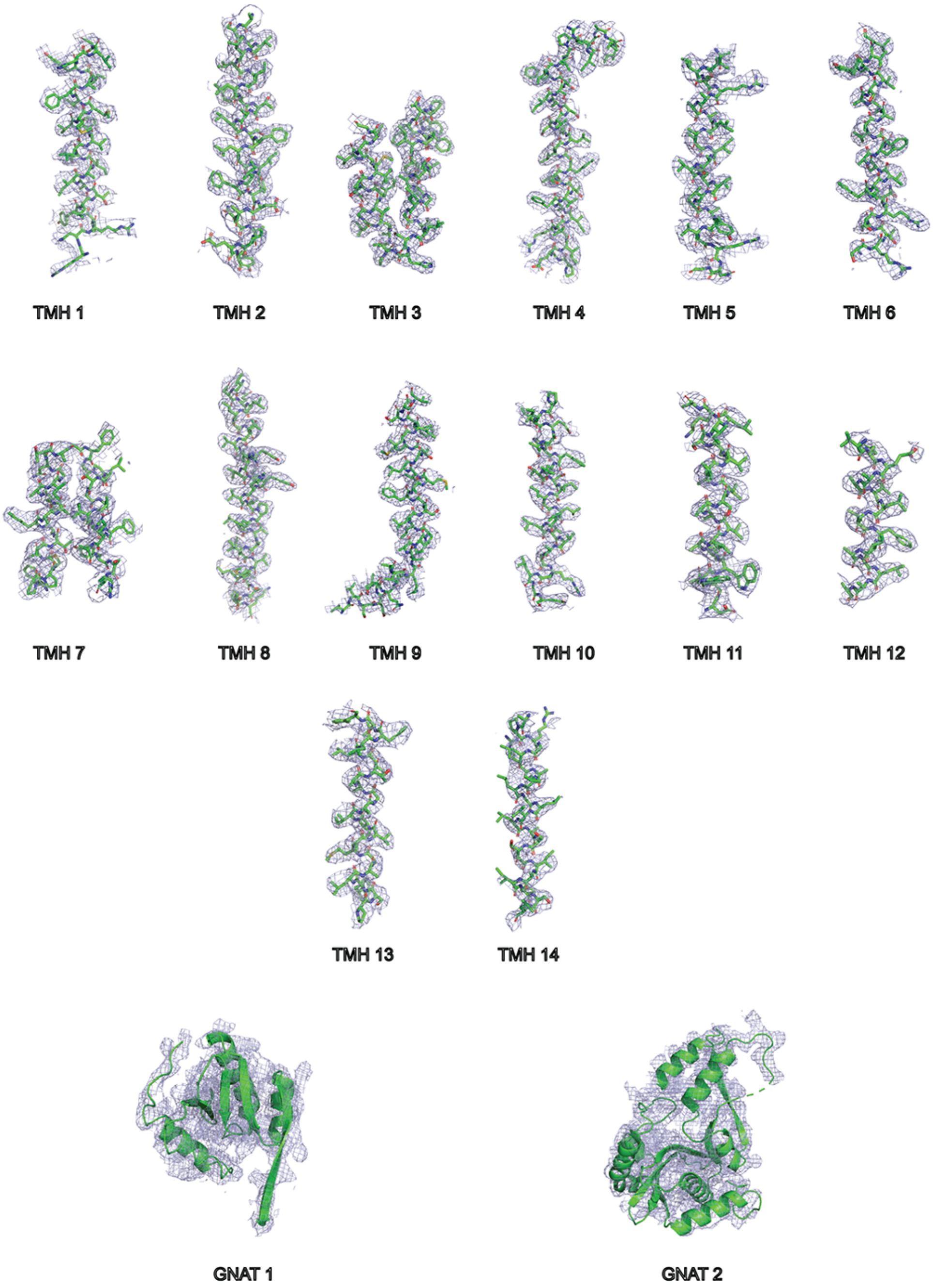


**Figure S4: Representative cryo-EM density of *AfMprF* in nanodisc.** The fit of the transmembrane helices and the soluble domain to the corresponding region of the final sharpened cryo-EM map (blue mesh) contoured at 6 σ. Figures were made with PyMoL[52].

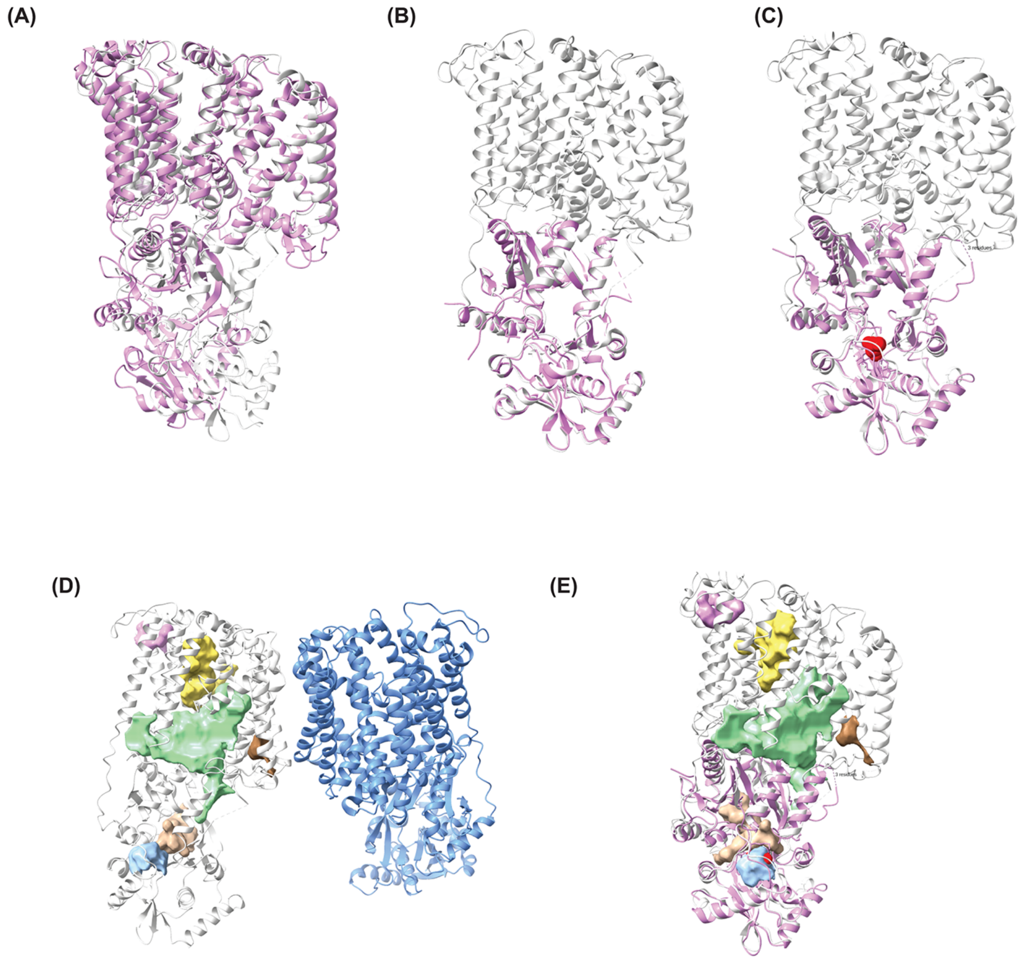


**Figure S5: A)** An overlay of the *AfMprF* monomer in gray with *Pa*MprF in violet (PDB – 9J8Q) with an RMSD of 4.25 Å (1.05 Å for 265 pruned atoms). Larger deviation is observed in the GNAT2 domain and the TM helices. Both models are derived from cryo-EM maps of the enzyme in GDN. **B)** An overlay of the full length *Af*MprF in gray with *Af*MprF synthase domain (violet) determined by x-ray crystallography (PDB-5VRV) with an RMSD of 0.73 Å. **C)** An overlay of the *AfMprF* in gray with *Bl*MprF synthase domain in violet (PDB – 4v36) with an RMSD of 1.1 Å. Lysinamide is shown as sphere (red). **D)** Potential cavities in full-length *Af*MprF analysed by KVfinder and a few cavities with >150 $Å^3$ are shown. The large cavities are found in the TM domain (green and yellow) on either side of the re-entrant helices. In the synthase domain, the two cavities highlighted here are possible substrate binding sites (blue and wheat). The monomers of *Af*MprF are colored in gray and blue. **E)** The synthase domain of *Bl*MprF (violet) with lysinamide is overlaid on *Af*MprF with the cavities highlighted. The lysnamide (red sphere, also in panel B) position matches the cavity in blue.
